# Eye movements reflect attentional navigation of number space without visual experience

**DOI:** 10.64898/2026.09.16.751961

**Authors:** Giuliano Giari, Federica Sigismondi, Roberto Bottini

## Abstract

Primates use eye movements to explore and gather information about their external, visual world. Similarly, even when there is nothing to see spontaneous eye movements reflect the structure and the exploration of internal, conceptual knowledge. However, it remains unclear what mechanism links eye movements to this internal search process. One possibility is that eye movements reflect visual imagery or visual exploration strategies used to explore conceptual knowledge. On the other hand, eye movements may reflect attentional movements in the conceptual space, which are independent from visual experience. Early blind individuals are a useful testbed for this hypothesis in that they never had access to functional vision, thus ruling out the influence of imagery or recycling of visual exploration strategies. To this end, we asked early blind individuals to perform a random number generation task while recording their eye movements. Using electrooculography (EOG), we found that eye movements track the exploration of the mental number line in both sighted and blind: participants moved their eyes to the right for positive changes in magnitude in the number sequence and to the left for negative changes, with eye movement amplitude scaling with the magnitude difference. Furthermore, sighted and early blind participants showed a similar temporal profile, with the effect peaking around 750 ms before number onset. These findings support the hypothesis that internally generated shifts of attention, rather than eye movements per se, drive the exploration of conceptual spaces.

**PUBLIC SIGNIFICANCE STATEMENT:** Spontaneous eye movements do not merely support visual exploration, but also reflect the exploration of internally represented conceptual spaces, such as numbers or colors. However, it remains unclear whether these eye movements depend on visual experience or whether they instead reflect an internal attentional mechanism. We tested early blind individuals and sighted participants in a random number generation task. We found that eye movements systematically reflected navigation through numerical space in both groups. Our results suggest that attention may serve as a general mechanism for moving through relational knowledge, with the oculomotor system revealing these internal dynamics even in the absence of visual experience.

## INTRODUCTION

Humans and non-human primates rely on eye movements to extract information from their surroundings (Nau, Julian, et al., 2018; Piccardi et al., 2016). Through sequences of saccades and fixations, eye movements support the construction of spatial relationships in the observer’s mind (Bicanski & Burgess, 2019; Lakshminarasimhan et al., 2020). The extracted relational knowledge is then stored in cognitive maps, whose activity is mainly supported by the hippocampal-entorhinal system (Bellmund et al., 2018; Buzsáki & Moser, 2013; Doeller et al., 2010; Moser et al., 2008; Nau, Navarro Schröder, et al., 2018; Rowland et al., 2016; Meister & Buffalo, 2016; Rolls & Wirth, 2018). Interestingly, navigation of conceptual knowledge is associated both with the recruitment of hippocampal cognitive maps in a very similar way to what is observed in physical space (Constantinescu et al., 2016; Jasmin et al., 2025; Park, 2021; Viganò et al., 2023), as well as with spontaneous eye-movements that reflects the geometry of conceptual spaces. For instance, when people are asked to speak out loud numbers in the dark, in random order, they tend to look right before saying a number bigger than the previous one and left before saying a smaller one (Loetscher et al., 2010; Viganò et al., 2024). Similarly, when asked to generate colors instead of numbers, eye movements track the semantic distance between color words (Viganò et al., 2024), and when people (or monkeys) are asked to recall or recognize items belonging to different categories (even with arbitrary category boundaries) eye movements partition the (empty or dark) visual space reflecting the learned categorical space (Rosen & Freedman, 2025; Caron & Ester, 2026). Yet, it remains unclear what underlies eye movements in conceptual spaces such as those described above. One possibility is that visual experience plays a crucial role in the development of this behavioral correlate, with individuals relying on visual imagery to navigate conceptual space by mentally reconstructing visual representations of relations between concepts, or simply repurposing strategies learned during visual behavior. Alternatively, eye movements in conceptual spaces may reflect movements of attention that are independent of visual experience and visual imagery, possibly reflecting a mental search in which attention moves from one item to the other, internally, similar to the external allocation of attention in the world (e.g., to visual, auditory or other sensory signals). Here, we aimed to disentangle the origins of eye movements during navigation of conceptual spaces by testing early blind individuals (who have never used vision functionally to move within a space and do not have any reported visual memory; see supplementary table 1), and using a numerical conceptual space as a testbed (Loetscher et al., 2010; Viganò et al., 2024). If eye movements during conceptual navigation are related to visual imagery or the reconstruction of visual representations of conceptual relationships, early blind individuals should not express this behavioral correlate during navigation of conceptual spaces. On the contrary, if they reflect shifts of attention, eye-movements congruent with a left-to-right mental number line should be observable independently of visual experience. To this end, we asked twenty-eight sighted and seventeen early blind individuals to perform a random number generation task in a darkened room, with no perceptible visual cues, while eye movements were recorded using electrooculography (EOG) and, in sighted, also with an infrared eye tracker. Participants were instructed to verbally generate numbers between 1 and 12 as randomly as possible, producing approximately one number every two seconds (see Methods, Fig1A). The experiment comprised 5 runs of 13 blocks each, with a 0.5 Hz metronome presented only during the first block of each run as a reminder of the pace participants were asked to keep while saying the numbers (see Methods).

## MATERIALS & METHODS

### Participants

A total of 51 participants were analysed for this study. Of these, 28 were sighted individuals (SC) collected as part of a larger investigation that included an MEG recording session in which we also recorded eye tracking and electroculogram (EOG) data. Only the eye tracking and EOG were used in the current manuscript. Similarly, 13 early blind individuals (EB) participated in an MEG recording session with concurrent EOG recordings that were analyzed in this manuscript. Another sample of 10 EB took part in an EOG-only recording session. In total, we analyzed the data of 28 SC and 23 EB.

Participants reported no history of neurological disorders. Prior to each session they gave written informed consent to participate in the experiment. All procedures were approved by the ethical committee of the University of Trento.

### Experimental design

The experimental design was the same in all recording sessions. Participants were asked to generate numbers (in Italian) in the range from 1 to 12 in the most random order possible (Viganò et al., 2024). Participants performed 5 runs of 13 30-second blocks in which they were asked to produce a number every 2 seconds, for a total of 14 numbers per block. Before the experiment participants were familiarized with the task by practicing 12 blocks with a metronome, following Loetscher et al., 2010 and Viganò et al., 2024. As compared to these other studies we opted for a slightly lower frequency of 0.5 Hz to allow for more time between words, necessary to carry out MEG analyses that are not part of the current manuscript. Moreover, we decided to exclude the metronome from most of the main experimental blocks in order to remove unnecessary neural responses due to the regular sound (i.e., frequency tagging). However, we kept the metronome in the first block of each run to help participants remember the speaking pace. Each block started and ended with an alerting sound at 1200 Hz. The metronome instead was an individual sound set at 800 Hz. The SC participants, during the 30s blocks were visually presented with static noise patterns (i.e., 0.4deg visual angle grayscale boxes covering the whole screen). This was done to promote eye movements (Salvaggio et al., 2019; Sahan et al., 2022).

Stimuli presentation, as well as the recording of participants’ voices, was controlled via MATLAB and Psychtoolbox.

### Audio recording and preprocessing

Audio traces were recorded separately for each 30s block. For the audio recording we used a microphone placed near the mouth of the participants of the EOG-only sample, while for the MEG sample a microphone was placed on the wall of the MEG shielded room to avoid interference with the MEG signals. Audio recordings were automatically segmented and transcribed using WhisperX (Bain et al., 2023). Transcriptions were then automatically checked for: i) words ending after the end sound; ii) presence of words different from numbers or nans; iii) duration of individual words exceeding 3 standard deviations of the block mean duration; iv) durations longer than 2 s; v) time difference between transcriptions being less than 1 s; vi) silence periods of more than 3 seconds; vii) repeated labels, to check for model’s hallucinations. If any of these criteria were present, the whole audio recording for that block was opened in Audacity and manually corrected by the experimenters (GG; FS). We then excluded from all subsequent analyses: i) numbers that were out of the 1-12 range; ii) words ending after the end sound.

#### Eye tracking and EOG data acquisition and preprocessing

### EOG-only sample

Unipolar EOG recordings were acquired at the Center for Mind/Brain Sciences of the University of Trento with a 8-channel DC system (Brain Products GmbH, software: BrainVision Recorder version 1.21) in an electromagnetically shielded booth. Continuous data were recorded at 1000 Hz. Horizontal EOG was placed at the right and left outer canthi. Reference and ground electrodes were placed along the forehead midline, with the former positioned in the upper forehead, close to the hairline, and the latter in the lower forehead, close to the eyebrows. We computed bipolar derivatives offline by taking the difference between channel pairs.

### MEG sample

We collected bipolar EOG data as auxiliary channels of the Elekta Neuromag 306 MEG system (Elekta, Helsinki, Finland) placed in an electromagnetically shielded room (AK3B, Vakuumschmelze, Hanau, Germany) at the Center for Mind/Brain Sciences of the University of Trento. Continuous EOG data were recorded at 1000 Hz with hardware bandpass filters in the range 0.1–330 Hz. For the SC participants, eye tracking data were also recorded continuously and streamed as additional channels in the MEG system. The eye tracker (Eyelink 1000 Plus, SR Research Ltd., Ottawa, Canada) was used to track the left eye. Before each run SC participants performed a standard 9-point calibration. Noise patterns were projected at 120 Hz on a translucent whiteboard positioned 1 m in front of the participant using a ProPixx projector (Vpixx Technologies, Canada).

Data analyses were carried out in python using MNE-python (Gramfort et al., 2014) and common scientific python tools (pandas, scipy, numpy). Continuous recordings from both samples were segmented based on words onsets obtained from the audio segmentation. Following previous work (Vigano et al., 2024, Loetscher et al., 2010), we focused on the time period preceding word onsets based on the hypothesis that this time window is informative about the process of searching in conceptual spaces. Specifically, we segmented the recordings from -1 to 0 s with respect to word onsets. Block-level timestamps were realigned to the continuous recordings using the triggers sent to the recording devices at the beginning of each block.

#### Correlation analyses

### Eye tracking-EOG correlation

We estimated the Pearson correlation between horizontal EOG and eye tracking recording on the x axis for each subject of the SC group. This analysis was done as a first sanity check of the reliability of EOG data to capture meaningful eye movements information.

### Correlation with numerical distances

We defined numerical distances as the signed difference between consecutively pronounced numbers within a 30 s block. This difference was computed before trial exclusion to ensure that it reflects the actual mental exploration process as experienced by the participant. Following Loetscher et al., 2010 and Vigano et al., 2024, we then computed the median eye position (either using eye tracking or EOG) in the 1 s window before word onset. For this analysis we excluded trials as defined earlier, plus the trials in which the same number was repeated. We then computed the Spearman correlation between eye position and numerical distances. Correlation scores were computed separately for each 30 s block, resulting in 65 scores per subject, that were averaged obtaining one correlation score per subject. This correlation analysis was also repeated over time, computing one correlation score per block at every time point between -1 and 0 before word onset, allowing us to estimate the evolution of the effect over time.

#### Statistical analyses

### Behavior

We computed descriptive statistics of participants’ number generation performance, such as: i) total amount of words produced; ii) amount of words produced for each number; iii) amount of words produced for each run. We then compared these metrics between groups, using: a two-sample t-test to compare the total amount of words produced; a linear mixed model with the formula “word_count ∼ group * run + (1| subject)” to compare the number of words produced per run; a Poisson generalized estimating equations model with the following formula “number_count ∼ group * number” to compare word counts for each number between groups.

### Correlation between EOG and eye tracking

Correlation scores between the horizontal EOG and eye tracking data were fisher transformed and entered into a one-sample t-test against zero.

### Eye movements

The average correlation scores of each subject were fisher transformed. Depending on whether the data were normally distributed, as assessed using the Shapiro–Wilk test, statistical significance was evaluated using either a paired-samples *t*-test or a Wilcoxon signed-rank test. The correlation scores over time were also fisher transformed and entered into a cluster permutation test (Maris & Oostenveld, 20017), to correct for multiple comparisons across time. In brief, the cluster permutation test computes a t-test at each time point. The time points surviving an uncorrected threshold of p<0.05 were aggregated to form clusters based on temporal contiguity and their sum t-score retained as the cluster score. This procedure was repeated 5000 times and the cluster with the highest aggregated t-score at each iteration retained to form a null distribution. The observed cluster score is then compared to the null distribution to compute a cluster p-value. We defined as significant clusters surviving a threshold of p<0.05.

## RESULTS

First, we investigated whether sighted and early blind participants were comparable in terms of task performance by computing the total number of words produced and the frequency of each number across groups.

The results showed that the two groups produced a comparable number of words both in each run considered separately (linear mixed model; main effect of group: β = −3.141, p = 0.54; main effect of run: β = −0.453, p = 0.6; group × run: β = 0.846, p = 0.48 ; Fig. 1C) and across the entire experiment (two-sample t-test: t(43) = −0.16, p = 0.87; Fig. 1D). Moreover, the groups did not differ in how frequently each number was generated (generalized estimating equations model; group effect: z = −0.237, p = 0.812; Fig. 1E). Thus, both groups showed comparable behavioral performance, suggesting that any differences observed in the eye-movement data would have been unlikely driven by task-related factors. We then move to eye-movement data and we use eye-tracker data from sighted individuals, aiming to replicate the number–eye correlation reported in previous studies (Loetscher et al., 2010; Viganò et al., 2024). For each participant, we computed a signed difference vector for the generated numbers by subtracting each number from the preceding one, with negative values indicating increases and positive values indicating decreases. For the eye-movement data, we computed the mean horizontal eye position during the 1-second interval preceding each verbal response and derived a corresponding signed difference vector (Fig. 1B), where negative values indicated leftward shifts and positive values rightward shifts. Correlating the two vectors revealed a significant positive association, consistent with previous reports (Loetscher et al., 2010; Viganò et al., 2024) (one sample t-test ; t(27) = 7.43, p < 0.001, Fig. 2A). To characterize the temporal dynamics of this effect, we examined the time course within the 1-second window (see Methods), revealing a sustained positive correlation that formed a significant cluster spanning −1000 to 0 ms prior to verbal onset (p < 0.001, cluster corrected, a = 0.05, Fig 2B).

**Figure 1.**
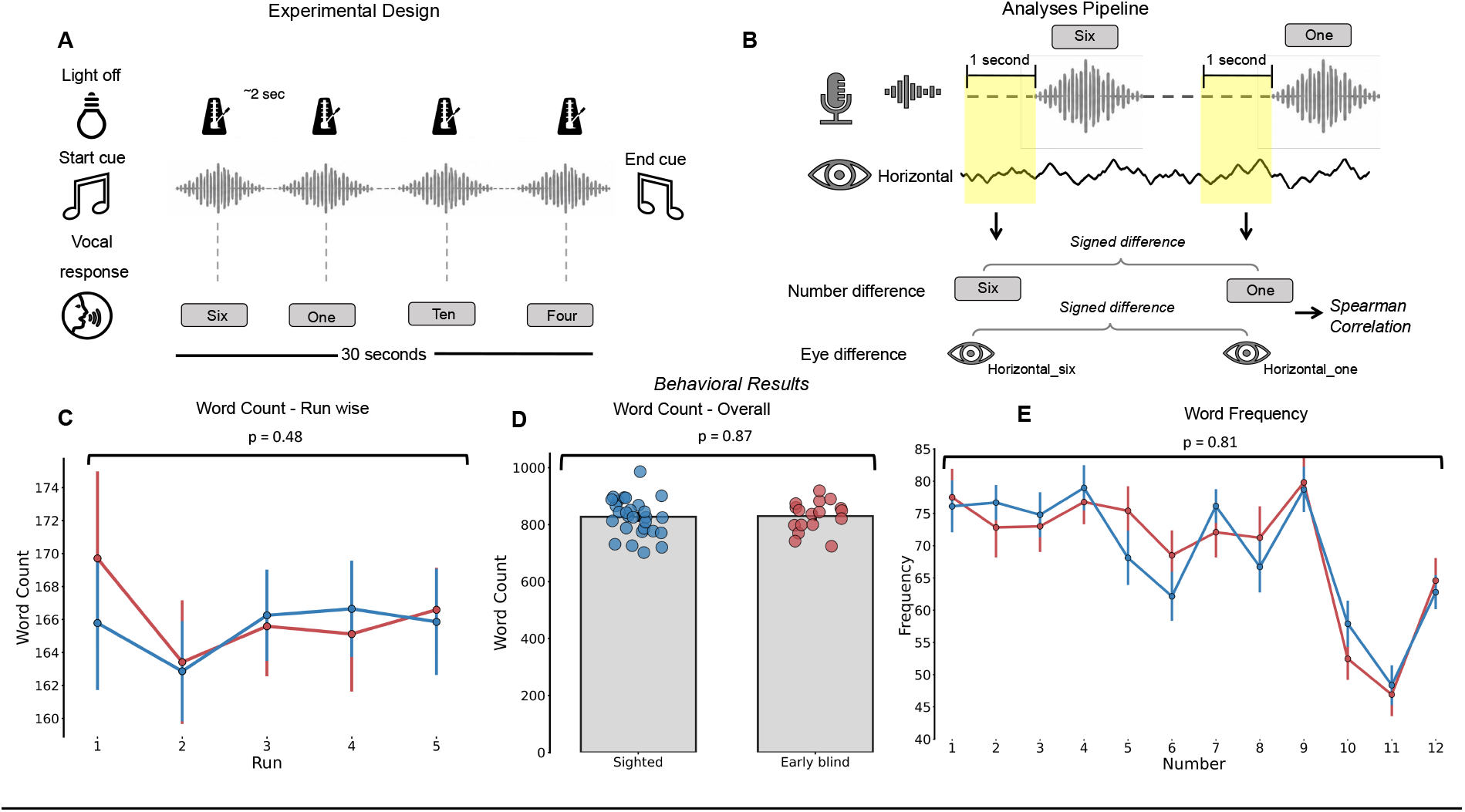
Experimental design and behavioral results. **A:** Schematic representation of the experimental design. Twenty-eight sighted and 17 early blind participants were asked to randomly generate numbers from 1 to 12 during 30-s blocks, at a fixed pace of approximately one number every 2 s. The task was performed in a “looking-at-nothing” paradigm, in which no salient visual information was presented on the screen, while eye position was recorded using electrooculography (EOG) in all participants and eye tracking in sighted participants only. The experiment consisted of five runs of 13 blocks each. During the first block of each run, a metronome set at 0.5 Hz was presented to help participants remember the required speech rate. **B:** Eye-position analysis pipeline. In sighted participants, for each produced number, we considered a 1-s time window preceding word onset and computed the median eye position within this interval. We then calculated the difference in eye position between consecutive number productions and related it to the corresponding numerical difference between the consecutively produced numbers. The signed difference in eye position was correlated with the signed numerical difference between consecutive numbers. The same analytical rationale was applied to early blind participants, but eye position was quantified within a ±200-ms time window centered on the peak of the effect identified in the early blind group (see Methods).. **C–D:** The number of words produced by sighted and early blind participants was comparable both when each run was considered separately (left; group × run interaction: β = 0.846, p = 0.48) and when considering the total number of words produced across the entire experiment (right; two-sample t-test: t(43) = −0.16, p = 0.87). **E:** The frequency with which each number was produced across the experiment did not differ between sighted and early blind participants (generalized estimating equations model; main effect of group: z = −0.237, p = 0.812).

**Figure 2.**
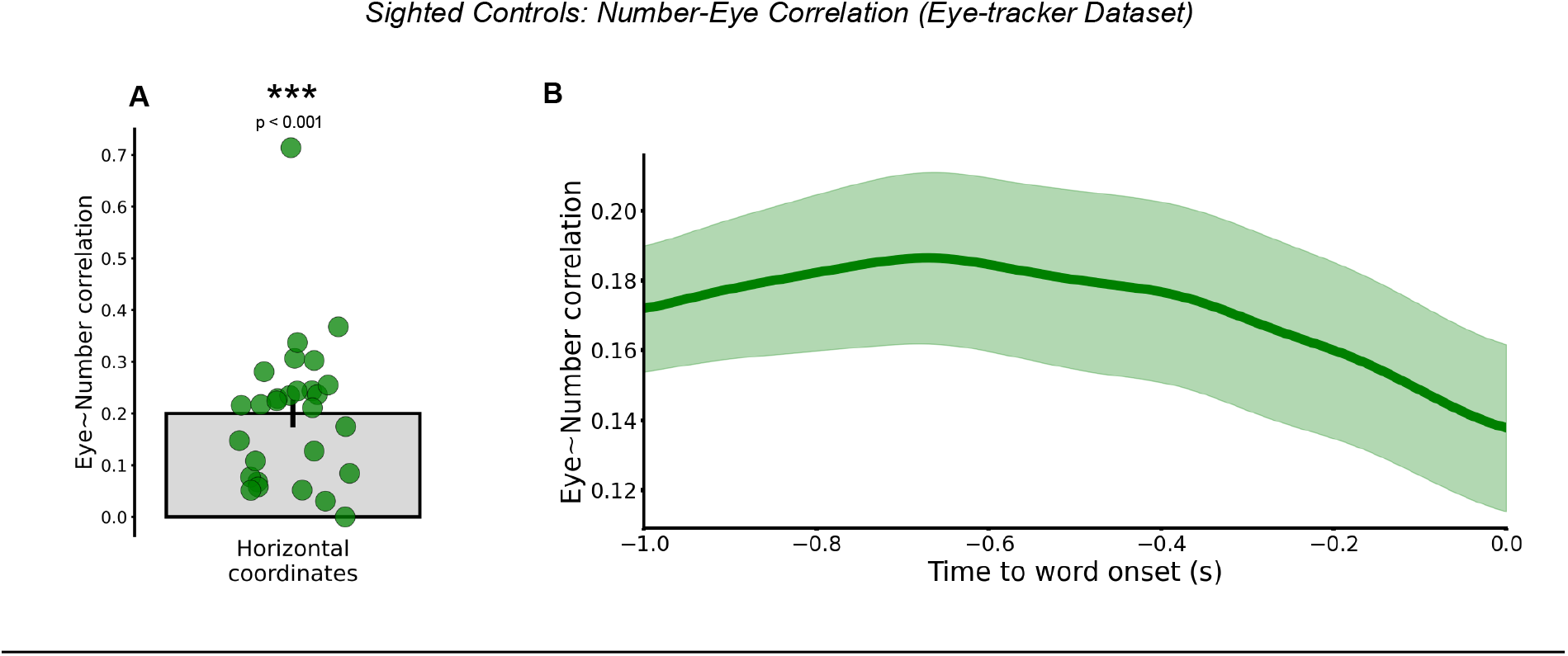
Eye–number correlations in sighted participants measured with eye tracking. **A–B:** Analysis of eye-tracker recordings from sighted participants revealed a significant correlation between the signed difference in eye position and the signed numerical difference between consecutive number productions (left; One sample t-test ; t(27) = 7.43, p < 0.001). This relationship indicates that eye movements preceding number production reflected the direction of navigation within numerical space: when participants were about to produce a number smaller than the preceding one, their gaze tended to shift leftward, whereas production of a larger number was associated with a rightward gaze shift. Inspection of the time course showed that this correlation was significant throughout the entire time window of interest (right; p < 0.001, cluster corrected).

This pattern suggests that the search process unfolds well before overt number production. Then we verify whether EOG could be a reliable method to capture gaze shifts also in a looking-at-nothing paradigm, where the signal could be noisier due to the absence of a reference screen or fixation point. To this end, we correlated the continuous horizontal (x-axis) eye-tracker time series with the corresponding horizontal EOG signal in the sighted group, observing a significant correlation between the two (mean r= 0.2; one sample t-test; t(27) = 4.84, p < 0.001; See Appendix B, Figure S1A–B). Together, these findings confirm that our experimental design replicates the previously reported coupling between eye movements and conceptual number navigation, and that EOG provides a reliable estimate of gaze position even in the absence of visual input. We then proceeded to analyze the eye movements of sighted and early blind individuals during the task as measured with EOG. In line with what was observed in the eye-tracker data, in sighted individuals we observed a significant positive correlation between the signed difference of the produced numbers and the signed difference of eye position (Wilcoxon test; W = 19, p < 0.001; Fig. 3A-B). Time-course analyses, once again, revealed a sustained positive correlation between the signed difference vectors (number and eye) across most of the time window, forming a significant cluster spanning −1000 to −200 ms, with a peak at -768ms, prior to verbal onset (p = 0.0002, cluster corrected, α = 0.05; Fig. 3C) and a gradual decay as speech onset approached. Similarly, early blind individuals showed a significant positive correlation between the signed difference of the produced numbers and the difference of eye position when considering the whole 1-s time window (Wilcoxon test; W = 31, p = 0.03, α = 0.05; Fig. 3D-E). We next examined the temporal dynamics of the effect in early blind individuals within the same 1-s time window. This analysis revealed two significant clusters, spanning −944 to −841 ms and −818 to −717 ms relative to number-production onset (p = 0.022 and p = 0.026, respectively; cluster-corrected, α = 0.05; Fig. 3F), and showing a peak of the effect (-759ms) comparable to the one found in the sighted. Comparing the effect between sighted and blind over this time window, we found only a trend toward significance (t(43) = 1.81, p = 0.08, α = 0.05; Fig. 3G), which was confirmed in temporally resolved analysis, with no cluster surviving correction for multiple comparison (all cluster-corrected p > .05, α = 0.05; Fig. 3H). Overall, these results support the hypothesis that eye movements reflect navigation along a mental number line in both sighted and early blind individuals. The two groups showed a similar temporal profile and comparable peak times.

**Figure 3.**
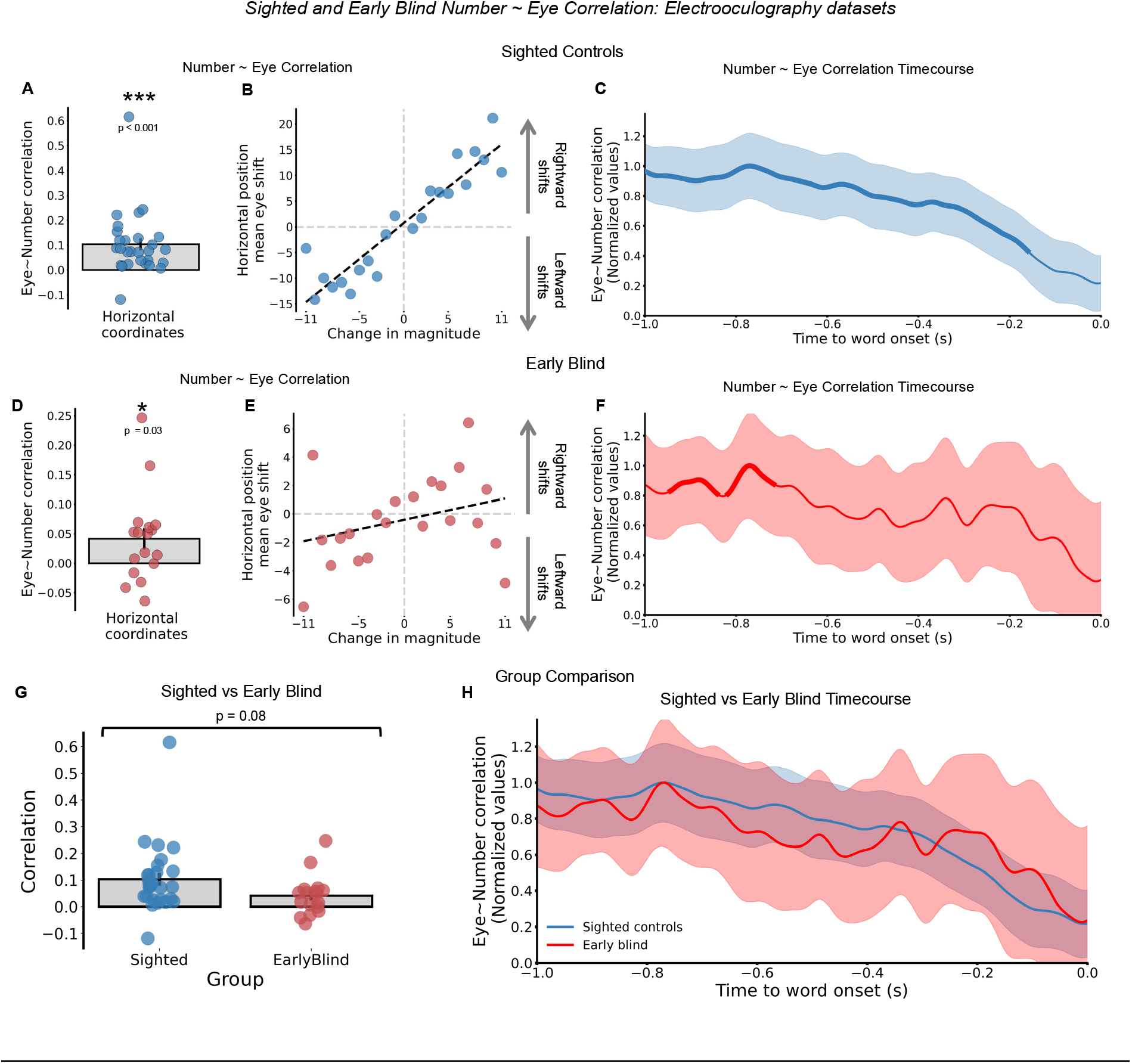
Eye movements in sighted and early blind individuals reflect navigation within an abstract number space. **A–B:** Analysis of electrooculography (EOG) data in sighted participants revealed a significant correlation between the signed difference in eye position prior to two consecutive word onsets and the signed difference between the two consecutively produced numbers (A, Wilcoxon test: W = 19, p < 0.001). This indicates that sighted participants tended to look more to the right when they were about to produce a number larger than the one previously said, and more to the left when they were about to produce a smaller number (B provides a visualization of this relationship for illustrative purposes only). **C:** Time-course analysis showed that this correlation was sustained from 1 s before word onset until 200 ms before word onset (p = 0.0002, cluster-corrected. Bold lines indicate significant clusters; The time course was smoothed for visualization purposes only). **D–E**: Similar to sighted participants, early blind participants analyses on EOG signal, also showed a significant correlation between signed eye-position difference and signed numerical difference (D, one-sample t-test: t(16) = 3, p = 0.007). This finding indicates that, even in individuals deprived of visual input from early in life, eye movements correspond to movements within an abstract number space. Panel E shows this relationship for visualization purposes only. **F:** Time-course analysis in early blind participants revealed two significant clusters of correlation: one spanning from −944 to −841 ms and a second from −818 to −717 ms relative to word onset, with a peak at −759 ms (p = 0.022 and p = 0.026, respectively; cluster-corrected. Bold lines indicate significant clusters, and the dashed line marks the peak of the effect. The time course was smoothed for visualization purposes only). **G–H**, The magnitude of the effect showed only a trend toward a significant difference between the two groups (two-sample t-test: t(42) = 1.82, p = 0.08). Time-resolved analyses, however, did not identify any significant clusters indicating a difference between sighted and early blind participants after correction for multiple comparisons (all p > 0.05).

## DISCUSSION

In this study, we tested sighted and early blind individuals in a random number generation task with the aim of characterizing the role of eye movements during navigation of abstract conceptual knowledge. We observed that in both sighted and early blind individuals, eye movements correlate with the change in magnitude between subsequent numbers. Indeed, in both groups, a higher magnitude difference between two consecutive numbers was associated with a rightward shift in eye position, whereas a smaller magnitude difference was associated with a leftward shift in eye position. Interestingly, both groups showed a very similar time course, with the peak of the effect around 770ms before the spoken word.

Several previous studies have shown that eye movements accompany the retrieval and exploration of internally represented information in a structured rather than random manner. In “looking-at-nothing” paradigms, eye position predicts the magnitude of the next number to be reported, suggesting that participants search along a spatially organized mental number line (Loetscher et al., 2010; Viganò et al., 2024). Similar effects have been observed for other conceptual domains: spontaneous eye movements reflect the representational geometry of color space (Viganò et al., 2024), and during memory retrieval people spontaneously direct their gaze toward distinct regions of empty space depending on the conceptual category they are retrieving (Viganò, Cristoforetti et al., 2025). Related findings further suggest that gaze may reflect the internal organization of behaviorally relevant categories, even when these categories are arbitrary and learned during the task (Rosen & Freedman, 2025; Caron & Ester, 2026). Together, these findings indicate that eye movements can provide an overt behavioral marker of internal search through structured representational spaces.

A recent theoretical account has proposed that conceptual navigation may rely on internally generated attentional movements, in analogy with bodily movements during physical navigation and gaze movements during visual exploration (Bottini et al., 2026). Within this framework, attention functions as an abstract form of action: shifts of attention provide a translation signal that allows cognitive maps to track the current position within a relational space. In primates and humans, hippocampal and entorhinal activity is modulated by gaze position and saccade direction during visual exploration (Kilian et al., 2015; Jutras et al., 2013; Georges-François et al., 1999), and entorhinal grid-like codes have been reported during the exploration of visual space (Staudigl et al., 2018; Kilian et al., 2012; Nau et al., 2018; Julian et al., 2018). Importantly, spatial codes in the hippocampal–entorhinal system can also be modulated by covert attention, even in the absence of overt eye movements (Wilming et al., 2018; Giari et al., 2023), and a recent paper has indeed found a correlation between eye movements during conceptual operations over a modified mental number line, and the representations of such operations in the entorhinal cortex (Eperon et al. 2026). These findings suggest a close relationship between gaze, attention, and the neural mechanisms supporting navigation through both physical and abstract spaces.

However, an important question remained unresolved: do eye movements during conceptual navigation reflect a mechanism learned through visual experience, such as an internalized form of visual exploration or visual imagery, or do they instead reveal a more general attentional mechanism that is independent of vision? Early blind individuals provide a critical test case for this question. Because they have never had access to functional vision, and typically report no visual memories, the presence of systematic eye movements during conceptual search would be difficult to explain in terms of visual imagery or recycled visual scanpaths.

Thus, our findings suggest that visually learned exploration strategies are not necessary for eye movements to track navigation through conceptual space. Rather, the most parsimonious interpretation is that eye movements provide an overt signature of internally generated attentional shifts within a structured representational space. In the present task, each produced number may act as a transient landmark within the mental number line, anchoring the search for the next candidate number. The observed gaze pattern then reflects the direction and distance of this internal attentional shift.

More broadly, our results support the idea that attention may serve as a general mechanism for moving through relational knowledge (Bottini et al. 2026). Eye movements, in this view, are not the cause of conceptual navigation, but a behavioral trace of the attentional dynamics through which internally represented spaces are explored. Importantly, this does not imply that early blind individuals do not engage in non-visual sensorimotor simulations, or even in non-visual imagery, during random number generation. Rather, it shows that, even if such simulations are involved, they are accompanied by attentional movements that engage the oculomotor system, likely because of its close coupling with the attentional system in the primate brain (Awh et al., 2006; Corbetta et al., 1998; Nobre et al., 1997).

To sum up, with this experiment we provide evidence that the eye movements observed during the navigation of conceptual spaces are not attributable to repurposed visual strategies or to visual imagery, but rather to movements of attention independent of visual experience.

## CONTSTRAINT ON GENERALITY

Our sighted and blind participants were all Italians, thus their mental representation of the number line is influenced by the occidental culture. The direction of the effect, i.e., left for large negative transitions and right for large positive transitions, may thus not generalize to other cultures. For instance, people with a different reading direction may instead show the opposite direction with large positive transitions being to the left. Furthermore, we tested only a specific case of spatial structure in the form of a one-dimensional mental number line. It remains to be tested whether these effects are found also in conceptual spaces of higher dimensionality. Nevertheless, there is no a priori hypothesis on why this should not be the case.

## ACKNOWLEDGEMENTS

We would like to thank the blind people who participated in the study. This work was supported by an ERC-CoG, ATCOM, grant n. 101125658 awarded to RB.

## AUTHORS CONTRIBUTIONS

Conceptualization: GG, FS, RB; Data curation: FS; Formal analysis: GG, FS; Funding acquisition: RB; Investigation: GG, FS; Methodology: GG, FS; Project administration: RB; Resources: GG, FS; Software: FS; Supervision: RB; Validation: FS; Visualization: FS; Writing – original draft: GG, FS, RB; Writing – review & editing: GG, FS, RB;

## DATA AND CODE AVAILABILITY

All data and analysis code will be made publicly available in appropriate repositories upon acceptance of the manuscript.

## APPENDIX A

**Supplementary table 1:**
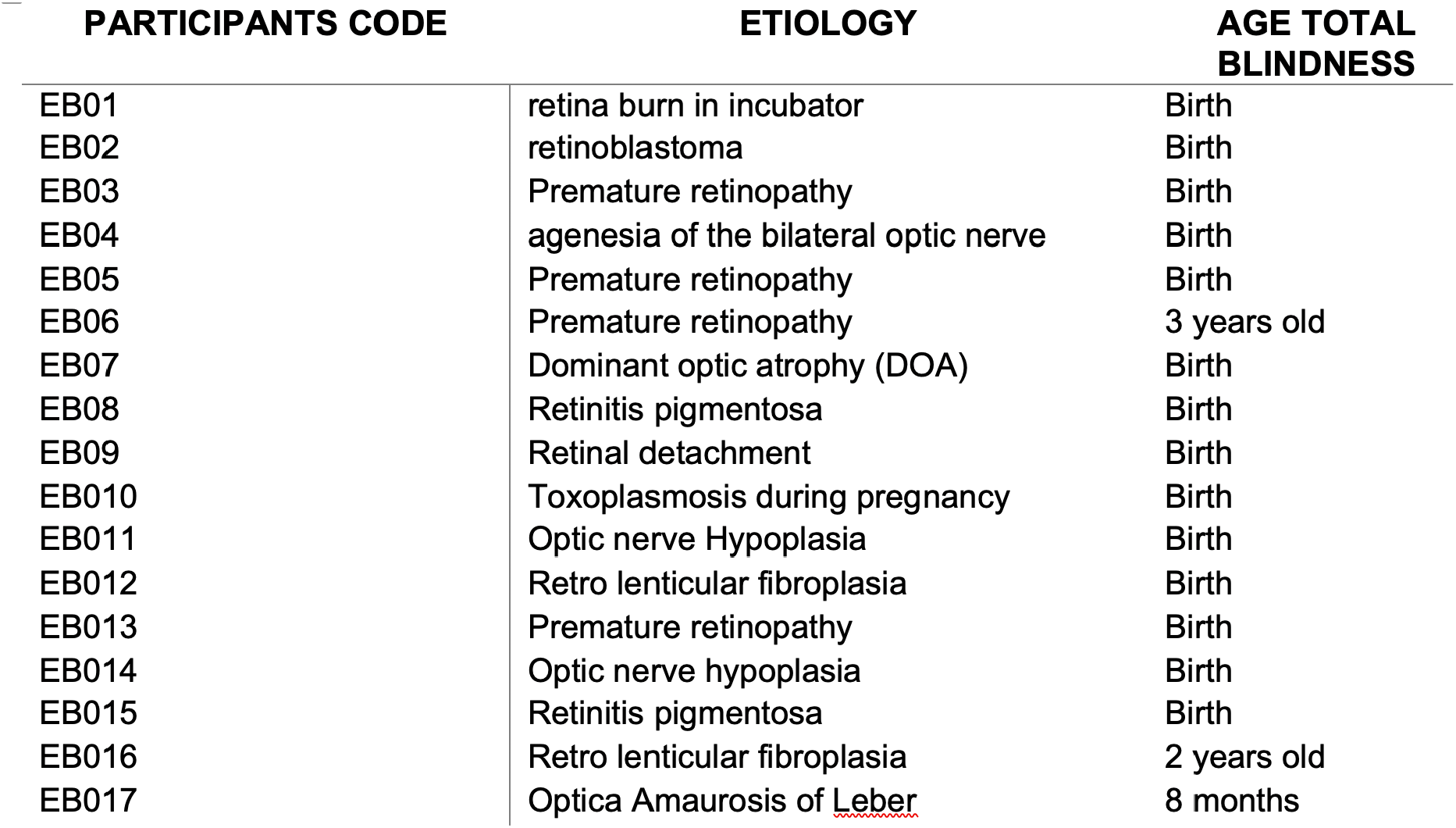
Early Blind Demographics.

| PARTICIPANTS CODE | ETIOLOGY | AGE TOTAL<br>BLINDNESS |
| --- | --- | --- |
| EB01 | retina burn in incubator | Birth |
| EB02 | retinoblastoma | Birth |
| EB03 | Premature retinopathy | Birth |
| EB04 | agenesia of the bilateral optic nerve | Birth |
| EB05 | Premature retinopathy | Birth |
| EB06 | Premature retinopathy | 3 years old |
| EB07 | Dominant optic atrophy (DOA) | Birth |
| EB08 | Retinitis pigmentosa | Birth |
| EB09 | Retinal detachment | Birth |
| EB010 | Toxoplasmosis during pregnancy | Birth |
| EB011 | Optic nerve Hypoplasia | Birth |
| EB012 | Retro lenticular fibroplasia | Birth |
| EB013 | Premature retinopathy | Birth |
| EB014 | Optic nerve hypoplasia | Birth |
| EB015 | Retinitis pigmentosa | Birth |
| EB016 | Retro lenticular fibroplasia | 2 years old |
| EB017 | Optica Amaurosis of <u>Leber</u> | 8 months |

## APPENDIX B

**Figure S1.**
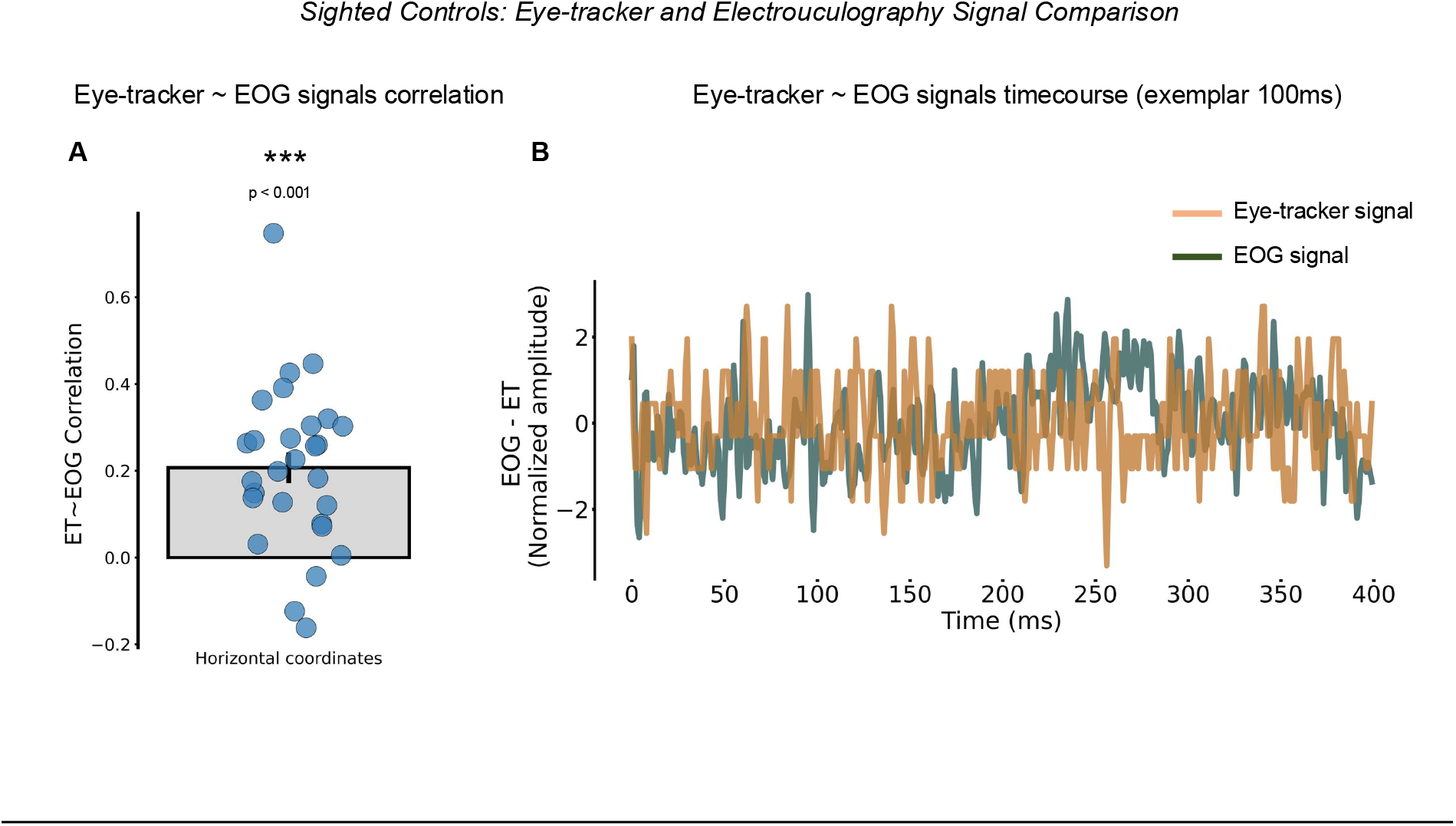
Eye-tracker and electrooculography signals provide comparable estimates of eye position.**A–B:** To verify that electrooculography (EOG) could be reliably used to estimate eye position under the experimental recording conditions, we correlated the eye-tracker signal obtained from sighted participants with the corresponding EOG signal and observed a significant correlation between the two (left; one sample t-test; t(27) = 4.84, p < 0.001). This indicates that the two measures captured comparable changes in eye position over time. Consistent with this, the overlaid signal traces showed a similar temporal profile (right; representative 100-ms segment shown for visualization purposes only).

